# A Comparison of Cell-Based and Cell-Cultured as Appropriate Common or Usual Names to Label Products Made from the Cells of Fish

**DOI:** 10.1101/2021.02.26.433119

**Authors:** William K. Hallman, William K. Hallman

**Author notes:** **Contact information for Corresponding Author:** Department of Human Ecology, 55 Dudley Rd. New Brunswick, NJ 08553.

## Abstract

Using an online experiment with a nationally representative sample of 1200 adult American consumers, two “common or usual names,” “Cell-Based Seafood” and “Cell-Cultured Seafood,” were assessed using five criteria. Displayed on packages of frozen Atlantic Salmon, the names were evaluated on their ability to differentiate the novel products from conventionally-produced fish, to identify their potential allergenicity, and after learning its meaning, to be seen by participants as an appropriate term for describing the process for creating the product. In addition, the names were evaluated as to whether they would be interpreted as disparaging of new or existing products, and whether they elicited reactions contrary to the assertion that the products are nutritious, healthy and safe. The results confirmed earlier research showing that “Cell-Based Seafood” slightly outperformed “Cell-Cultured Seafood” as a common or usual name. Labeling products with the term “Cell-Based Seafood” meets important regulatory criteria by enabling consumers to distinguish such products from conventional seafood products, and by indicating the presence of allergens. From a marketing perspective, “Cell-Based” is also viewed as an appropriate term for describing the process for producing the products, meeting the criteria for transparency. Consumers also had more positive reactions to “Cell-Based Seafood” and were slightly more inclined to want to taste and purchase “Cell-Based” products both before and after learning the meaning of “Cell-Based” and “Cell-Cultured.” Therefore, “Cell-Based Seafood” should be adopted as the best common or usual name to label cell-based seafood products.

**Practical Application:** Widespread adoption and consistent use of a single “common or usual name” for “Cell-Based” seafood, meat, poultry and other products by the food industry, regulators, journalists, marketers, environmental, consumer, and animal rights advocates, and other key stakeholders would help shape public perceptions and understanding of this rapidly advancing technology and its products. This study confirms that “Cell-Based Seafood” is the best performing term to label seafood products made from the cells of fish. It meets relevant FDA regulatory requirements and slightly outperforms “Cell-Cultured Seafood” with regard to positive consumer perceptions, interest in tasting and likelihood of purchasing these novel products.

## 1 Introduction

Development of the technology to bring cell-based meats, poultry, and seafood to market at an affordable price is moving at a rapid pace (Dolgin, 2020; Miller, 2020). Stakeholder adoption and consistent use of a single term to refer to and to label cell-based protein products would help settle regulatory issues, shape public perceptions, and promote a clearer understanding of cell-based products (Hallman & Hallman, 2020). Yet, consensus regarding what to call these products still remains elusive, with different stakeholders favoring different terms (Ong, Choudhury, Naing, 2020).

Much of the research designed to answer this question of nomenclature has focused on issues of consumer acceptance of cell-based meat products (Bryant & Barnett, 2018, 2020). This approach makes sense from a marketing perspective since the promised benefits of cell-based meats, poultry, and seafood (Stephens et al., 2018; Tomiyama et al., 2020) can only be realized if consumers are willing to purchase them. However, the term ultimately used to label cell-based products must meet regulatory criteria as well as marketing criteria. Names chosen to maximize potential consumer acceptance (Szejda, 2018) may fall short of regulatory requirements or may be viewed as false or misleading by regulators. U.S. Food and Drug Administration (FDA) regulations (21CFR101.3) call for foods that lack defined *standards of identity* (21CFR130.8) to be labeled with a *statement of identity*, such as a “common or usual name” to help inform consumer choices about food products available for purchase. Correspondingly, the US Department of Agriculture (USDA) requires that meat (9CFR317.2) and poultry products (9CFR381.117) be labeled using common or usual names. The FDA and the USDA Food Safety and Inspection Service (USDA-FSIS) have formally agreed to jointly regulate cell-based meat and poultry (though seafood would be regulated solely by the FDA) (Post et al., 2020; U.S. Food and Drug Administration and U.S. Department of Agriculture Office of Food Safety, 2019).

Key to common or usual names under 21CFR102.5 is that the specified name simply, directly and accurately describe or identify the basic nature of the food or the ingredients or properties that distinguish it from other products. It also must not be easily confused with the name of another food that is not in the same category, and it should convey what the product is in a clear way that differentiates it from other foods.

Balancing both marketing and regulatory considerations, Hallman and Hallman (2020) proposed five criteria for choosing a common or usual name that could be used to appropriately label products made from the cells of fish, shellfish, and crustaceans, and by extension, other cell-based meat, poultry, and game products. In their criterion A, they argued that to meet FDA and USDA regulatory requirements, a common or usual name should enable consumers to distinguish cell-based products from conventionally produced products. For seafood, this means that the common or usual name should signal to consumers that the cell-based seafood is neither wild-caught nor the product of aquaculture (i.e., farm-raised).

While Hallman and Hallman’s criterion A is that the common or usual name convey that there are important differences between cell-based and conventional products, their criterion B is that the common or usual name should also signal important similarities. FALCPA, the Food Allergen Labeling and Consumer Protection Act of 2004 (Public Law 108-282) requires that foods that consist of, or that contain protein from a “major food allergen,” bear a label that declares that allergen’s presence. Because cell-based seafood products will necessarily be produced using the cells of fish, shellfish, or crustaceans, the common or usual name should not suggest that the products are safe to eat by those who are allergic to other seafood products.

While meeting FDA regulatory requirements is a necessary prerequisite, the common or usual name must also meet the needs of consumers and the companies making these products. While perhaps implicit in the FDA requirements for common or usual names, Hallman and Hallman (2020) set as their Criterion E, that consumers view the name as appropriate to identify the product. Consumers increasingly demand transparency in food labeling (FMI and Label Insight, 2020). Moreover, because of the purported environmental, ethical and other benefits associated with cell-based meat, poultry, and seafood, companies should want to transparently differentiate their cell-based products from their conventional counterparts. They may also find such differentiation necessary to justify the price premium likely needed to be charged when cell-based products initially make it to market. In choosing to voluntarily differentiate their products using a transparent common or usual name, producers of cell-based meat, poultry, and seafood would also likely preempt efforts to mandate labeling of their products using terms they may find limiting or pejorative

Finally, producers of cell-based meat will want to avoid repeating the errors made in introducing GM (genetically modified) foods to consumers. One of the mistakes made by producers of GM foods was to send unlabeled GM products into Europe and other markets where they faced significant resistance. The resulting backlash created longstanding mistrust of producers of GM products and of GMOs in general (Mohorčich & Reese, 2019).

Hallman and Hallman (2020) also argued that a common or usual name should be chosen that is not viewed as “disparaging” of either existing conventional products or cell-based products (Criterion C). Similarly, they suggest that an effective common or usual name should not elicit consumer reactions that suggest that the cell-based food products are unsafe, unhealthy, or less than nutritious (Criterion D). These latter criteria recognize that if the common or usual name is expected to be adopted voluntarily by producers, it cannot work against efforts to sell either cell-based or conventional products. Producers of cell-based products have already rejected terms proposed by some consumer organizations (Hansen, 2018) such as “lab-grown meat,” “synthetic meat,” “artificial meat,” and “fake meat. Producers assert that these terms are scientifically inaccurate and are intended to portray their foods as artificial and unpalatable (AMPS Innovation, 2020). At the same time, traditional meat producers have rejected names they believe are disparaging of their own conventional products. These include names preferred by animal rights advocates and some companies, including “clean meat,” “animal-free meat,” “slaughter-free meat,” and “cruelty-free meat” (Greene & Angadjivand, 2018).

Hallman and Hallman (2020) used these five criteria as the basis for testing seven potential common or usual names for cell-based seafood. The names they tested included “Cultivated Seafood,” “Cultured Seafood,” “Cell-Based Seafood,” and “Cell-Cultured Seafood.” They also tested the phrase, “Produced using Cellular Aquaculture,” and the phrases “Cultivated from the Cells of ____,” and “Grown directly from the Cells of ____,” filling in the blanks with the name of the packaged seafood product. Three controls (wild-caught, farm-raised, and no common or usual name) were also tested as comparisons. To test these names and phrases, they used a 3 × 10 between-subjects experimental design, collecting data online from a quota sample of 3,186 US adults drawn from opt-in panels. These common or usual names tested were shown as labels on realistic packages of frozen seafood (salmon, shrimp and tuna).

The results showed that all of the common or usual names performed equally well in signaling that those allergic to seafood should not eat the products (Criterion B). Each was also seen as an appropriate name to identify the product (Criterion E).

However, the majority of consumers were unable to differentiate seafood products labeled with the terms “Cultivated,” “Cultured,” and the phrase “Produced using Cellular Aquaculture” from conventional “Wild-Caught” or “Farm-Raised” seafood. In fact, 54% of those who saw the term “cultivated,” 41% of those who saw the term “Cultured,” and 39% of those who saw the phrase “Produced using Cellular Aquaculture” wrongly assumed that the products were “Farm-Raised.” Therefore, none of these terms meet the essential regulatory criterion (A) for common or usual names. Only the four terms incorporating the word “cell” (“Cell-Based,” “Cell-Cultured,” “Cultivated from the Cells of ____,” and “Grown directly from the Cells of ____”) cued more than half of the participants that the products were neither “Wild-Caught” nor “Farm-Raised.”

However, the phrases “Cultivated from the Cells of ____” and “Grown directly from the Cells of ____” performed poorly with respect to the consumer perception / marketing criteria. Consumers rated products with those terms the least positively and they were seen as most likely to be genetically modified. Importantly, they also performed relatively poorly regarding consumer perceptions of the associated product’s taste, safety, nutrition, and naturalness, particularly in comparison to conventional “Wild-Caught” and “Farm-Raised” products. Consumers also expressed the least interest in tasting, and were least likely to purchase the products with these terms.

Both of the names, “Cell-Based” and “cell-cultured,” signaled to more than half of the participants that the product differs from both “Wild-Caught” and “Farm-Raised” seafood (meeting criterion A). In direct comparisons, the terms “Cell-Based” and “Cell-Cultured” were not significantly different from each other on most of the consumer perception and marketing related measures tested. Nevertheless, “Cell-Based” was found to outperform “Cell-Cultured” when comparing the pattern of results for each term to those of the conventional “Wild-Caught” and “Farm-Raised” seafood products, with which these novel products would compete in the marketplace. Therefore, Hallman and Hallman (2020) concluded that the term “Cell-Based” was the better name.

While Hallman and Hallman (2020) recommended “Cell-Based” as the best performing term of the seven tested, “Cell-Based” and “Cell-Cultured” generated similar results. The study also had some limitations. It was designed as an initial evaluation of seven potential common or usual names (and three comparisons) and tested these using three different seafood products. The resulting 3 × 10 experimental design randomly assigned ~100 participants per condition. Because no statistically significant interactions were found between the common or usual name tested and the type of seafood product, tests of main effects of common or usual name were able to be conducted with samples of ~300 per condition. This provided sufficient power to detect relatively small differences in means and proportions among the 10 names in the analysis. However, because of the large number of statistical tests performed, conservative p-values needed to be adopted to reduce experiment-wise error. In addition, the opt-in quota sample of ~300 per condition is inadequate to project the results to the US population with a reasonable margin of sampling error.

To overcome these limitations, this study examines the two best performing names identified by Hallman and Hallman (2020), “Cell-Based” and “Cell-Cultured,” using a nationally representative sample of 1200 participants, permitting projections of the study results to the population. It also adds additional measures to further explore consumer perceptions of the nature of the products, and their perceptions of the products after learning the meaning of the common or usual names.

Many consumers are likely to first encounter these novel products through seeing a package in a grocery store. Therefore, common or usual names must convey meaning on their own—that is, without additional explanation on the label. Following the eventual regulatory clearance and introduction of the products into the marketplace and with the adoption and use of a consistent common or usual name, consumer awareness, knowledge, and understanding of the products and the technology used to produce them will likely grow over time. This study therefore also adds measures of consumer perceptions of the products *after* reading an explanation of the meaning of the terms.

## 2 Materials and Methods

### 2.1 Experimental Design

Two proposed common or usual names, “Cell-Based Seafood” and “Cell-Cultured Seafood” were tested. Each participant was randomly assigned to view only one of the names, which were tested on the labels of high-definition images of packages of frozen Atlantic Salmon Fillets. Salmon was chosen because it is one of the most often consumed seafood products in the U.S., so many consumers are familiar with it (Seafoodhealthfacts.org, 2018). Consistent with this, Hallman and Hallman (2020), found that 58.4% of their participants had eaten salmon in the previous year and that those assigned to view a salmon product were moderately familiar with salmon in general. Salmon is also high in Omega 3 fatty acids and low in methylmercury, so it is recommended by the FDA and EPA as a “best choice” for consumption by women who are (or might become) pregnant, breastfeeding mothers, and young children (U.S. Food and Drug Administration, 2019).

### 2.2 Materials

High-resolution pictures of the front of packages containing frozen Atlantic Salmon were created for this experiment, identical to those used in Hallman and Hallman (2020) (see Figure 1). These were designed to mimic conventional seafood packages currently available in the supermarket. As is typical of such packages, the top one-third depicted a cooked salmon fillet, presented as a “serving suggestion.” The middle third displayed the product title, “Atlantic Salmon Fillets.” The common or usual name to be tested was printed directly below the product title. A Nutrition Facts Label (NFL) with accurate values corresponding to those of conventional Atlantic Salmon Fillets appeared on the bottom third of the package. The net weight was printed at the bottom of the package along with declarations that the product “CONTAINS SALMON,” and is “PERISHABLE,” and advising consumer to “KEEP FROZEN” and to “COOK THOROUGHLY.”

**Figure 1.**
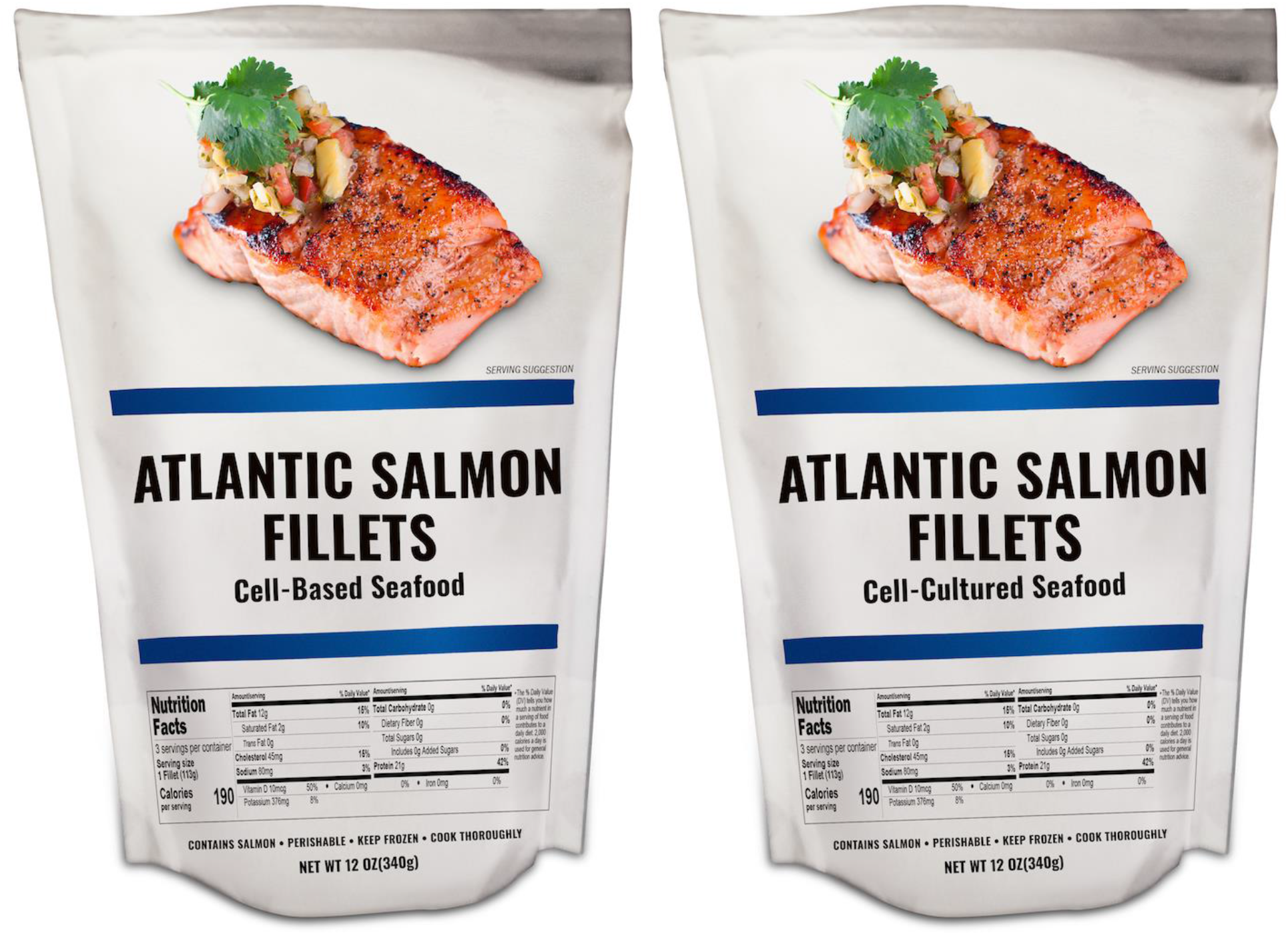
Package Images.

### 2.3 Participants

Data was collected between October 6 and October 13, 2020. The study participants consisted of adult American consumers (18 and older) recruited from the YouGov.com web-based consumer panel. YouGov initially interviewed 1780 respondents from whom, a sample of 1600 participants were selected to produce the final dataset, matching a sampling frame derived from the 2018 American Community Survey (ACS).

Of these 1600 participants, 1200 were randomly assigned to complete one of the two experimental conditions reported in this study, while 400 participants completed a related task to be summarized in a separate article. Through random assignment, a total of 591 participants viewed packages displaying the common or usual name, “Cell-Based Seafood,” while 609 viewed packages displaying the common or usual name, “Cell-Cultured Seafood.” Sampling error associated with N=600 is +/− 4% when projected to the population.

### 2.4 Procedure

The procedures used were adapted from those reported in Hallman and Hallman, 2020. The participants provided informed consent and confirmed that they were 18 years of age or older and so eligible to participate. They then read an inclusive description of the term “seafood” and were asked how often they had eaten a meal containing seafood in the previous 12 months, and if they had not eaten any seafood to indicate why. Those who had consumed seafood were then shown a list of seafood and asked to indicate which products they had eaten. The participants were also asked about their familiarity with dietary guidelines for eating seafood, and how many four-ounce portions of seafood they had eaten in the prior week.

The participants were then shown the image of the package bearing the common or usual name they had been randomly assigned. The participants were asked to look at the package carefully, to record (in free text) the “first thought, image, or feeling that comes to mind when seeing this package,” and then to rate how positive or negative this response was.

To ensure that each participant actively considered the package and its label, the participants saw the package a second time and were asked to repeat the same exercise. Finally, they were presented with the package a third time and asked how positive or negative their overall reactions to the salmon product were, how interested they would be in tasting the salmon, and if it were sold in their grocery store, how likely they would be to purchase it in the next six months.

The participants then viewed an enlarged version of the picture of the cooked salmon fillet that appeared on the package. They were then asked how familiar they are with salmon overall, whether they had ever tasted Atlantic Salmon, and if so, how much they liked or disliked the taste. Those who indicated that they had previously eaten salmon were asked if they had ever ordered a salmon fillet in a restaurant, purchased it in a store, online, or at a fish market. They were also asked about their likelihood to purchase uncooked and fully-cooked salmon fillets in a store in the next six months, whether they have ever cooked salmon fillets, whether it is true or false that salmon is a good source of “heart-healthy” Omega 3s, and if they, or anyone who lives in their households is allergic to salmon or to any other seafood.

The participants were then shown an enlarged image of the product name “Atlantic Salmon Fillets” along with the common or usual name to be tested printed below it. While viewing the image, the participants were asked, “Which of the following best describes this salmon?” The response categories were “Wild-Caught,” “Farm-Raised,” and “Neither Wild-Caught nor Farm-Raised.” Those who indicated that it was “Neither Wild-Caught nor Farm-Raised” were then asked a follow-up question, “Which of the following best describes this salmon?” with the response categories, “Made from the cells of Salmon,” “Made from the cells of Plants,” and “Made from neither Salmon nor Plant cells.”

Participants were asked whether those allergic to fish should eat the salmon, as well as how safe it would be to consume the salmon if one is not allergic to fish. They then rated the product’s naturalness and how likely they thought that it had been genetically modified.

The Nutrition Facts Label (NFL) was then shown, enlarged so that it could be easily read. While the NFL was still on screen, the participants indicated how nutritious the salmon is, and how good or bad they thought the salmon tastes. Finally, they were asked whether pregnant women should eat the salmon and separately, whether children should consume it.

Because a common or usual name must convey appropriate meaning on its own, no definition of either “Cell-Based” or “Cell-Cultured” Seafood was provided to the participants prior to the final part of the experiment. Participants then read the following description (“Cell-Cultured Seafood” was substituted for those randomly assigned to that condition).

> “The term <u>Cell-Based Seafood</u> indicates that this salmon differs from both wild-caught and farmed salmon. It tastes, looks, and cooks the same and has the same nutritious qualities as Atlantic Salmon produced in traditional ways. Yet, it involves a new way of producing just the parts of salmon that people eat, instead of catching or raising them whole. <u>Cell-Based Seafood</u> means that a small number of cells from Atlantic Salmon were placed in a nutrient solution, where they grew and reproduced many times. The resulting meat was then formed into fillets that can be cooked or eaten raw.”

After reading this definition, the participants were asked to indicate their existing familiarity with “the *idea* of producing just the parts of salmon that people eat, instead of catching or raising them whole.” They were asked to indicate how appropriate the term was “for describing this new way of producing just the parts of salmon that people eat, instead of catching or raising them whole?” They then rated the clarity of the term in communicating that the product “was not caught in the ocean,” how clear it communicated that the product was not farm-raised, and whether they agreed or disagreed that Atlantic Salmon that is “Cell-Based” (or “Cell-Cultured”) should be “sold in the same section of the supermarket as wild-caught and farm-raised fish.”

After having read the description of “Cell-Based” (or “Cell-Cultured”) Seafood, the participants were prompted to take a final look at the package of Atlantic Salmon. They were then asked how positive or negative their overall reactions to the salmon were, how interested they would be in tasting it, how likely they would be to buy the product in the next six months if it were sold in their grocery store, and how likely they would be to recommend that pregnant women buy the salmon. They then answered questions related to a second experiment, the results of which will be summarized in a subsequent article. The participants finished by reporting whether they have any children under the age of five living in the household and whether they are the primary shopper in their household.

### 2.5 Statistical Analyses

Analyses were conducted using IBM SPSS Statistics for Windows (version 27; IBM Corp., Armonk, New York). Differences in means were analyzed using Analysis of Variance to produce effect sizes using partial eta-squared (η_p_^2^). Z-tests of column proportions with Bonferroni correction were used to analyze differences in proportions. A p-value of 0.05 was used to distinguish significant differences within statistical tests. Where appropriate, weighted data is reported in the tables reporting percentages projected to the US population. To avoid potential distortions in the variance associated with key variables, sample weights were not used when reporting means, standard deviations, the results of ANOVAs, effect sizes, and correlations.

## 3 Results and Discussion

The median length of the experiment reported here was approximately 11.8 minutes. Consistent with census data, 51.3% of the 1200 participants were female. Mean age was 47.41, SD=17.69; 10.8% reported children under age 5 in the household. When asked “who does the grocery shopping for the household,” 55.4% reported doing “all of it,” 17.7% “most of it,” 15.5% “about half of it,” 8.5% “some of it,” and 2.9% “someone else does all of it.” Additional sociodemographic characteristics of the sample provided by YouGov as part of its panel recruitment are shown in Table 1.

**Table 1.** Sociodemographic Characteristics of the Sample, (N) = 1200

| Sociodemographic Characteristic* | % of total |
| --- | --- |
| Gender |  |
| Male | 48.7% |
| Female | 51.3% |
| Marital status |  |
| Married | 44.7% |
| Single, never married | 33.2% |
| Divorced or separated | 14.2% |
| Living with partner | 6.2% |
| Widowed | 5.8% |
| Educational level |  |
| Less than high school | 4.7% |
| High school /GED | 33.8% |
| Some college | 23.0% |
| 2-year college degree (Associate) | 8.7% |
| 4-year college degree (BA, BS) | 18.4% |
| Post-Graduate | 11.5% |
| Race/Ethnicity |  |
| White | 63.1% |
| Black/African-American | 12.1% |
| Hispanic/Latino | 16.2% |
| Asian | 3.5% |
| Native American | 1.3% |
| Two or More Races | 2.1% |
| Other | 1.6% |
| Middle Eastern | 0.2% |
| Household income |  |
| Less than \$10,000 | 6.8% |
| \$10,000 to \$19,999 | 8.5% |
| \$20,000 to \$29,999 | 12.9% |
| \$30,000 to \$39,999 | 11.1% |
| \$40,000 to \$49,999 | 7.7% |
| \$50,000 to \$59,999 | 6.9% |
| \$60,000 to \$69,999 | 6.0% |
| \$70,000 to \$79,999 | 8.3% |
| \$80,000 to \$119,999 | 4.2% |
| \$120,000 to \$249,999 | 1.8% |
| \$250,000 to \$349,999 | 1.7% |
| \$350,000 to \$499,999 | 0.6% |
| \$500,000 or more | 0.4% |
| Prefer not to say | 7.9% |

\*Categories and data provided by YouGov, collected as part of their panel recruitment.

About nine-in-ten (90.5%) of the participants reported having eaten one or more meals containing seafood in the 12 months prior to the survey. Moreover, 63.6% reported they had eaten at least one seafood meal a month, 31.4% reported that they had eaten at least one seafood meal a week, and 1.2% indicated that they had consumed one or more meal containing seafood per day. About four-in-ten (42.9%) reported having eaten a salmon fillet in the previous 12 months. Only 8.1% reported that they were “not familiar at all” with salmon in general. Consistent with this, 70.0% reported that they had previously purchased uncooked salmon fillets in a store, online, or at a fish market, 69.5% reported that they had cooked salmon fillets, and 42.0% reported that they had ordered a salmon fillet in a restaurant. The majority (58.6%) reported having previously tasted Atlantic Salmon specifically, with 83.5% of these indicating that they liked its taste.

The remaining results are structured to address the specific criteria described in the introduction.

### 3.1 Criterion A – Ability to distinguish from conventional products

A fundamental regulatory criterion for an acceptable common or usual name is its capacity to signal that the labeled product is different from those that consumers may already be familiar with. To test this, the participants were shown the product packages three times and asked to provide reactions to them. They were then asked, “Which of the following best describes this salmon?” Is it best described as “wild-caught,” “farm-raised,” and “neither wild-caught nor farm-raised”?

As shown in Table 2, the majority of those who viewed the name “Cell-Based” (60.1%) and those who saw “Cell-Cultured” (58.9%) on the package label correctly identified the salmon as “neither wild-caught nor farm-raised.” There were no statistically significant differences in these percentages, projected to the population. Thus, even in the absence of additional labeling information describing their meaning, both names do a good job of indicating to American consumers that the products are different from conventional wild-caught and farm-raised fish. However, a greater proportion of those who saw the name “Cell-Cultured” (30.1%) assumed that the product was farm-raised than those who saw the name “Cell-Based” (24.9%). In contrast, a greater proportion of those who saw the name “Cell-Based” (15.0%) assumed that the product was wild-caught than those who saw the name “Cell-Cultured” (11.1%).

**Table 2.**
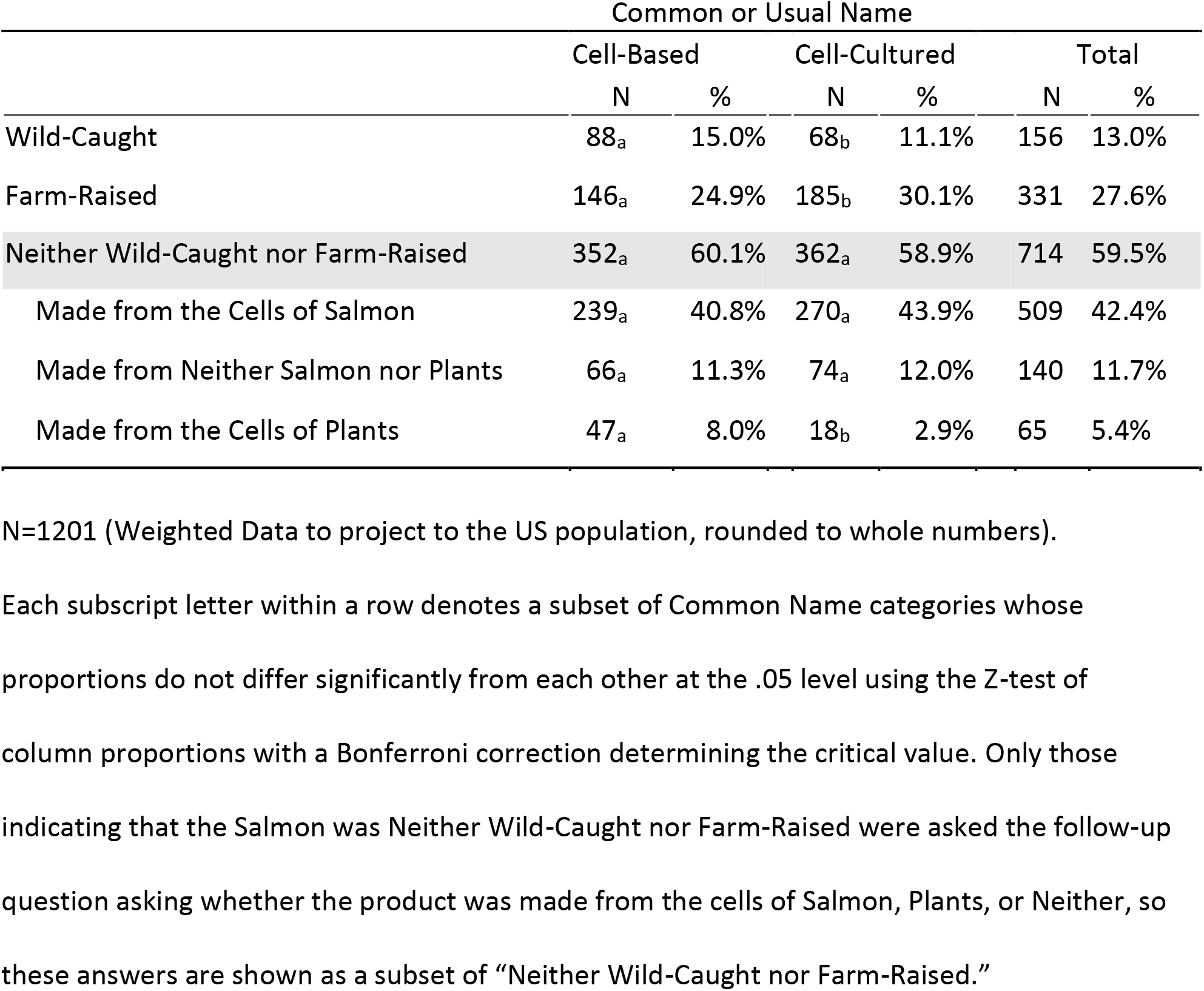

|  | Common or Usual Name |  |  |  |  |  |
| --- | --- | --- | --- | --- | --- | --- |
|  | Cell-Based |  | Cell-Cultured |  | Total |  |
|  | N | % | N | % | N | % |
| Wild-Caught | 88<sub>a</sub> | 15.0% | 68<sub>b</sub> | 11.1% | 156 | 13.0% |
| Farm-Raised | 146<sub>a</sub> | 24.9% | 185<sub>b</sub> | 30.1% | 331 | 27.6% |
| Neither Wild-Caught nor Farm-Raised | 352<sub>a</sub> | 60.1% | 362<sub>a</sub> | 58.9% | 714 | 59.5% |
| Made from the Cells of Salmon | 239<sub>a</sub> | 40.8% | 270<sub>a</sub> | 43.9% | 509 | 42.4% |
| Made from Neither Salmon nor Plants | 66<sub>a</sub> | 11.3% | 74<sub>a</sub> | 12.0% | 140 | 11.7% |
| Made from the Cells of Plants | 47<sub>a</sub> | 8.0% | 18<sub>b</sub> | 2.9% | 65 | 5.4% |

N=1201 (Weighted Data to project to the US population, rounded to whole numbers).
Each subscript letter within a row denotes a subset of Common Name categories whose
proportions do not differ significantly from each other at the .05 level using the Z-test of
column proportions with a Bonferroni correction determining the critical value. Only those
indicating that the Salmon was Neither Wild-Caught nor Farm-Raised were asked the follow-up
question asking whether the product was made from the cells of Salmon, Plants, or Neither, so
these answers are shown as a subset of “Neither Wild-Caught nor Farm-Raised.”

The participants who correctly responded that the salmon was “Neither wild-caught nor farm-raised,” were asked to indicate whether the salmon could be best described as “Made from the cells of Salmon,” “Made from the cells of Plants,” or “Made from neither Salmon nor Plant cells.” As shown in Table 2, the largest percentage of those who viewed “Cell-Cultured” (43.9%) and of those who viewed “Cell-Based” (40.8%) indicated that “Made from the cells of Salmon” was the best descriptor for the product. There are no statistically significant differences in these percentages, projected to the population. Thus, even in the absence of additional labeling, both names do a good job of indicating to American consumers that the products are made from the cells of fish. The smallest percentage (8.0%) of those who saw “Cell-Based” and “Cell-Cultured” (2.9%) thought that the product was “Made from the cells of Plants.” A z-test of column proportions indicated that these proportions are statistically different. A similar proportion (11.3%) of those who viewed “Cell-Based,” and 12.0% of those who viewed “Cell-Cultured” thought that the product was made from “neither plant nor salmon cells.”

### 3.2 Criterion B – Signal the presence of potential allergens

The proteins in the cells of fish can cause allergic responses in some individuals. Therefore, it is important that consumers recognize that cell-based seafood products will also contain potential allergens and avoid eating them. To test this, participants were shown the product title and common or usual name, and were asked, “If you are allergic to fish, is it safe for you to eat this salmon?” The response options were, 1 definitely not, 2 probably not, 3 probably yes, 4 definitely yes. “Cell-Based” and “Cell-Cultured” were equally competent in signaling allergenicity (*H*(1)=1.687, *p* =.194). Overall, participants understood that those with allergies to fish should *not* eat the product (*Mdn*=2.0).

### 3.3 Criteria C and D - Not be viewed as disparaging of cell-based or conventional products

The participants were asked to carefully examine the package of seafood shown to them and asked to type their response to the question, “What is the first thought, image, or feeling that comes to mind when seeing this package?” They were then asked to look at the package a second time and to record the thought, image, or feeling that came to mind. Each of the responses was coded using one of the 28 categories developed by Hallman and Hallman (2020) (see Table S1 in the supplemental materials). Each response was independently coded by two trained researchers, with any discrepancies resolved by consensus.

After recording their open-ended responses, each participant rated how positive or negative their thought, image, or feeling was, using a scale ranging from 1 extremely negative to 7 extremely positive. They were then asked to look at the package a third time and using the same scale, record how positive or negative their overall reaction was.

As shown in Table 3, the thoughts, images, and feelings associated with “Cell-Based” were rated by the participants as more positive than those associated with “Cell-Cultured.” Similarly, the participants’ overall reaction to “Cell-Based” was also rated more positively than their overall reaction to “Cell-Cultured.”

**Table 3.** Ratings of Thoughts, Images, or Feelings and Overall Reactions By Common or Usual Name

| | M | SD | N | F | P-value | $\eta^2$ |
| --- | --- | --- | --- | --- | --- | --- |
| Rating of First Thought, Image or Feeling |  |  |  | 10.267 | < 0.001 | .022 |
| Cell-Based | 4.84 | 1.78 | 591 |  |  |  |
| Cell-Cultured | 4.49 | 1.94 | 609 |  |  |  |
| Rating of Second Thought, Image or Feeling |  |  |  | 7.633 | < 0.01 | .018 |
| Cell-Based | 4.69 | 1.73 | 591 |  |  |  |
| Cell-Cultured | 4.40 | 1.91 | 609 |  |  |  |
| Overall Reactions |  |  |  | 11.514 | < 0.001 | .023 |
| Cell-Based | 4.82 | 1.72 | 591 |  |  |  |
| Cell-Cultured | 4.46 | 1.93 | 591 |  |  |  |

Scale: 1 extremely negative; 2 moderately negative; 3 slightly negative; 4 neither positive nor negative; 5 slightly positive; 6 moderately positive; 7 extremely positive.

The participants were asked how safe it would be to eat the salmon if one is not allergic to fish, responding using the scale: 1 very unsafe; 2 moderately unsafe; 3 somewhat unsafe; 4 neither safe nor unsafe; 5 somewhat safe; 6 moderately safe; 7 very safe. Both the “Cell-Based” (*M* = 5.58, *SD* = 1.64) and “Cell-Cultured” Salmon (*M* = 5.54, *SD* = 1.65) were equally rated as “somewhat” to “moderately” safe to eat (*F*(1, 1198) =0.178, *p* = .673, η_p_^2^ = .000). They were also equally rated as “moderately” nutritious; “Cell-Based” (*M* = 3.55, *SD* = 0.95), “Cell-Cultured” (*M* = 3.55, *SD* = 0.98), (*F*(1, 1197) = .002, *p* = .97, η_p_^2^ = .000) [Scale: 1 not at all nutritious; 2 slightly nutritious; 3 moderately nutritious; 4 very nutritious; 5 extremely nutritious].

Both products were also equally imagined to taste “slightly” good; “Cell-Based” (*M* = 5.09, *SD* = 1.59), “Cell-Cultured” (*M* = 4.99, *SD* = 1.64), (*F*(1, 1198) = 1.337, *p* = .25, η_p_^2^ = .001) [Scale: 1 extremely bad; 2 moderately bad; 3 slightly bad; 4 neither good nor bad; 5 slightly good; 6 moderately good; 7 extremely good]. The participants also reported that they were “moderately” interested in tasting both products, though they were slightly more interested in tasting “Cell-Based” (*M* = 3.12, *SD* = 1.49) than “Cell-Cultured” Atlantic Salmon (*M* = 2.94, *SD* = 1.52), (*F*(1, 1198) = 4.499, *p* = .034, η_p_^2^ = .004), [Scale: 1 not at all interested, 2 slightly interested, 3 moderately interested, 4 very interested, 5 extremely interested].

Both products were equally rated as “neither natural nor unnatural”; “Cell-Based” (*M* = 4.22, *SD* = 1.87) and “Cell-Cultured” Salmon (*M* = 4.07, *SD* = 1.96), (*F*(1, 1197) = 2.033, *p* = .154, η_p_^2^ = .002) [Scale: 1 very unnatural, 2 moderately unnatural, 3 somewhat unnatural, 4 neither natural nor unnatural, 5 somewhat natural, 6 moderately natural, 7 very natural]. However, “Cell-Cultured” Salmon (*M* = 5.62, *SD* = 1.43) was seen as slightly more likely to have been genetically modified than “Cell-Based” Salmon (*M* = 5.42, *SD* = 1.52), (*F*(1, 1198) = 5.395, *p* = .02, η_p_^2^ = .004) [1 extremely unlikely; 2 moderately unlikely; 3 slightly unlikely; 4 neither likely nor unlikely; 5 slightly likely; 6 moderately likely; 7 extremely likely].

Overall, the participants believed that pregnant women should probably not consume *either* of the salmon products. Using weighted data, 53.6% of the participants seeing either name indicated that pregnant women should probably or definitely not eat this salmon. Coded as 1 definitely not, 2 probably not, 3 probably yes, and 4 definitely yes, the median for both “Cell-Based” and “Cell-Cultured” was 2.00. By contrast, the majority in both conditions indicated that children *should* eat the salmon depicted using the same scale. The median for both “Cell-Based” and “Cell-Cultured” was 3.00. About seven-in-ten of those who saw “Cell-Based” (70.6%) and “Cell-Cultured” (69.1%) indicated that children should probably or definitely eat the salmon. Kruskal-Wallis tests indicated no statistically significant differences between the two names with respect to either dependent measure.

### 3.4 Criterion E – Be seen as an appropriate term

After viewing the description of the meaning behind “Cell-Based” or “Cell-Cultured,” two thirds of the participants (68%) reported that they were “not familiar at all” “with the *idea* of producing just the parts of seafood that people eat, instead of catching or raising them whole.” The remaining participants indicated that they were “slightly” (10.7%), “Moderately” (11.1%), “very” (6.5%) or “extremely familiar” (3.5%) with the idea (all percentages reported using weighted data). Coded on a scale of 1 not at all familiar to 5 extremely familiar, there were no statistically significant differences between the two names with regard to participant familiarity with the concept (*M* = 1.68, *SD* = 1.12). Similarly, using a scale of 1 “extremely inappropriate” to 7 “extremely appropriate,” both of the names were seen identically as “slightly appropriate” (*M*=4.97, *SD* = 1.81) “for describing this new way of producing just the parts of salmon that people eat, instead of catching or raising them whole.”

Participants were also asked how clear the term they viewed is, “in communicating that the salmon was not caught in the ocean,” and in communicating that it was not “Farm-Raised,” responding using the scale: 1 extremely unclear; 2 moderately unclear; 3 slightly unclear; 4 neither clear nor unclear; 5 slightly clear; 6 moderately clear; 7 extremely clear. The participants who saw “cell-cultured” indicated that the term was slightly clearer in communicating that, “the salmon was not caught in the ocean” (*M* = 4.52, *SD* = 2.07), than those who saw “Cell-Based” (*M* = 4.12, *SD* = 2.18), (*F*(1, 1198) = 10.48, *p* = .001, η_p_^2^ = .009). Similarly, “Cell-Cultured” was seen as slightly clearer in communicating that “the salmon was not farm-raised” (*M* = 4.38, *SD* = 2.09), than “Cell-Based” (*M* = 4.09, *SD* = 2.16), (*F*(1, 1198) = 5.315, *p* = .021, η_p_^2^ = .004).

It should be noted that these responses were given *after* reading the explanation of the meaning of the terms. Yet, when seeing the terms “Cell-Based” and “Cell-Cultured” on the packages at the beginning of the experiment (prior to explaining their meaning), both were seen equally as “Neither Wild Caught nor Farm Raised.” Moreover, a greater proportion of those who saw the name “Cell-Cultured” assumed that the product was farm-raised than those who saw the name “Cell-Based,” while a greater proportion of those who saw the name “Cell-Based” thought that the product was “Wild-Caught.” On its own, therefore, “Cell-Cultured” does not appear to be clearer than “Cell-Based” in demonstrating that the salmon was not produced using traditional methods.

The participants were asked to indicate their level of agreement that the “Cell-Based” and “Cell-Cultured” salmon they viewed should be sold in the same section of the supermarket as “Wild-Caught” and “Farm-Raised” seafood, using a scale of 1 strongly disagree to 7 strongly agree. The mean responses for both terms were identical, (*M*=4.31, *SD* = 1.90), [4 = “neither agree nor disagree”].

### 3.5 Consumer perceptions post-explanation of the meaning of the term

In the final part of the experiment the participants were prompted to take a final look at the package of salmon, and to consider it again, “now that you know what “Cell-Based” [or “Cell-Cultured”] means.” Repeating the same questions as those in the first part of the experiment, the participants were asked how positive or negative their reactions were to the salmon. The participants who saw packages labeled as “Cell-Based” had slightly more positive overall reactions (*M* = 4.24, *SD* = 1.93) than those who saw packages labeled as “Cell-Cultured” (*M* = 4.01, *SD* = 1.93), (*F*(1, 1198) = 4.164, *p* = .042, η_p_^2^ = .003) [Scale: 1 extremely negative to 7 extremely positive]. Those who saw “Cell-Based” also expressed slightly more interest in tasting the salmon (*M* = 2.83, *SD* = 1.47) than those who saw “Cell-Cultured” (*M* = 2.65, *SD* = 1.51), (*F*(1, 1198) = 4.397, *p* = .036, η_p_^2^ = .004) [Scale: 1 not interested at all to 5 extremely interested). Those who saw “Cell-Based” also indicated greater likelihood of purchasing the salmon in the next six months (*M* = 3.77, *SD* = 2.22) than those who saw “Cell-Cultured” (*M* = 3.45, *SD* = 2.26), (*F*(1, 1198) = 6.308, *p* = .012, η_p_^2^ = .005) [Scale: 1 extremely unlikely to 7 extremely likely). However, they were equally unlikely to recommend that pregnant women buy the salmon; “Cell-Based” (*M* = 3.34, *SD* = 1.97), “Cell-Cultured” (*M* = 3.26, *SD* = 2.03), (*F*(1, 1198) = 0.488, *p* = .485, η_p_^2^ = .000) [Scale: 1 extremely unlikely to 7 extremely likely).

### 3.6 Determining the best performing common or usual name

Each of the five criteria were assessed to determine the name which best meets the requirements of producers, consumers, and regulatory agencies. The results confirmed the original findings in Hallman and Hallman (2020). Nearly 80% of the participants indicated that were “not familiar at all” or only “slightly familiar,” “with the *idea* of producing just the parts of seafood that people eat, instead of catching or raising them whole.” Yet, on their own, both “Cell-Based Seafood” and “Cell-Cultured Seafood” signaled to 60% of consumers that the novel product differs from conventional “wild-caught” and “farm-raised” salmon (meeting criterion A) and without any additional explanation, more than 40% directly understood that the products were made from the cells of salmon. Both terms were equally able to signal potential allergenicity, with 72.6% of those who saw “Cell-Based Seafood” and 75.4% of those who saw “Cell-Cultured Seafood” indicating that those allergic to seafood should “probably” or “definitely not” consume the product (meeting criterion B) and both terms are seen as appropriately descriptive (meeting criterion E). Both are seen as equally safe and nutritious and are presumed to taste equally as good. Neither is seen as unnatural, although the products labeled as “Cell-Cultured” were seen as slightly more likely to have been genetically modified.

However, packages of Atlantic Salmon Fillets with the common or usual name “Cell-Based Seafood” were rated by participants as slightly more positive than those with the common or usual name “Cell-Cultured Seafood.” Both before and after reading the description of the meaning of the terms, participants reported more positive overall impressions, greater interest in tasting, and greater likelihood of purchasing the products labeled as “Cell-Based Seafood” than those labeled as “Cell-Cultured Seafood.”

It should be noted that the mean differences and associated effect sizes in these measures are quite small, though the pattern of those differences are consistent. These results also add to those of Hallman and Hallman (2020), who found that the pattern of results associated with “Cell-Based” were similar to those of “Wild-Caught” and “Farm-Raised” seafood products, while those associated with “Cell-Cultured” were dissimilar. In that study, initial reactions to “Cell-Based Seafood” were as positive as they were to both “Wild Caught Seafood” and “Farm Raised Seafood.” The products labeled as “Cell-Based Seafood” were also judged to be as nutritious as both “Wild-Caught” and “Farm-Raised” seafood, while “Cell-Cultured” products were not. Participants imagined that “Cell-Based Seafood” tasted as good as both “Wild-Caught” and “Farm-Raised” seafood. They were also equally interested in tasting and likely to purchase “Cell-Based Seafood” as they were seafood that was either “Wild-Caught” or “Farm-Raised.” In contrast, those who saw “Cell-Cultured Seafood” products were only as interested in tasting and purchasing them as they were in tasting and purchasing “Farm-Raised” seafood products.

Thus, the overall pattern of results from this study and that of Hallman and Hallman (2020) suggest that “Cell-Based” is the better choice for a common or usual name based on measures of likely consumer acceptance and purchase of these innovative products.

## 4 Conclusion

This study confirms that “Cell-Based Seafood” is the best candidate for a common or usual name for seafood made from the cells of fish. It meets the regulatory requirements to signal (on its own) that the novel products are not the same as conventional wild-caught and farm-raised seafood. At the same time, combined with the product name, “Atlantic Salmon Fillets,” it indicates to consumers that the products are made from the cells of fish, and therefore, those who are allergic to fish should not eat them. From a marketing perspective, “Cell-Based” is viewed as an appropriate term for describing the process for producing the products, meeting the need for transparency in labeling. Additionally, consumers indicate that they view “Cell-Based Seafood” products more positively than “Cell-Cultured” and are slightly more inclined to want to taste and purchase “Cell-Based” products. Therefore, the term “Cell-Based Seafood” should be considered the best common or usual name to be used to label seafood products produced from the cells of fish.

## Acknowledgments

This project was supported by BlueNalu.

## Author Contributions

W. Hallman is responsible for all aspects of the study, including its design, analysis of the data, and production of the manuscript. W. Hallman II assisted with the literature review, the coding of open-ended responses, and the review and final editing of the manuscript.

## Supplemental Material

**Table S1.** Open-Ended Thoughts, Images, and Feelings Categorized

|  | First Thought, Image or Feeling |  |  |  | Second Thought, Image or Feeling |  |  |
| --- | --- | --- | --- | --- | --- | --- | --- |
|  | Cell-Based | Cell-Cultured | Total |  | Cell-Based | Cell-Cultured | Total |
| None/IDK | 5.1% | 4.1% | 4.6% |  | 4.9% | 5.4% | 5.2% |
| Delicious/Appetizing/Yum/<br>Want to Eat/Try/Buy | 17.6% | 17.2% | 17.4% |  | 13.9% | 9.2% | 11.5% |
| Amazing/Awesome/<br>Attractive/Cool/Good/Great<br>/ Like it/Love it | 12.2% | 12.0% | 12.1% |  | 8.3% | 6.6% | 7.4% |
| Ok/ Acceptable | 0.8% | 0.3% | 0.6% |  | 2.0% | 1.1% | 1.6% |
| Bad/Disgusting/ Yuk<br>Unappetizing/Unappealing | 5.9% | 7.2% | 6.6% |  | 6.6% | 7.6% | 7.1% |
| Artificial/Fake/Not Natural/<br>Lab Grown/Manufactured | 3.2% | 6.6% | 4.9% |  | 3.6% | 3.4% | 3.5% |
| GMO | 0.0% | 0.8% | 0.4% |  | 0.2% | 0.3% | 0.3% |
| Concerned/Worried/<br>Unhealthy/Bad for you | 0.7% | 2.1% | 1.4% |  | 1.0% | 1.6% | 1.3% |
| Common Name Question | 12.2% | 8.5% | 10.3% |  | 8.3% | 9.4% | 8.8% |
| Common Name | 2.4% | 3.8% | 3.1% |  | 2.2% | 1.1% | 1.7% |
| Salmon | 6.1% | 3.9% | 5.0% |  | 3.6% | 3.6% | 3.6% |
| Salmon Preparation | 1.4% | 1.3% | 1.3% |  | 3.0% | 2.1% | 2.6% |
| Nutritional Aspects | 2.0% | 3.1% | 2.6% |  | 5.1% | 6.7% | 5.9% |
| Healthy/Good for You/Natural/Organic | 4.6% | 3.3% | 3.9% |  | 6.1% | 5.9% | 6.0% |
| Question/Confusion | 2.9% | 2.3% | 2.6% |  | 6.3% | 7.4% | 6.8% |
| Curious/Interesting | 2.4% | 1.5% | 1.9% |  | 0.7% | 1.8% | 1.3% |
| New/Innovative/Unfamiliar/Different | 0.1% | 0.6% | 0.4% |  | 1.2% | 0.7% | 0.9% |
| Do Not Like/Eat Fish/Salmon | 1.4% | 2.0% | 1.7% |  | 1.9% | 1.5% | 1.7% |
| Frozen/Not Fresh | 2.4% | 1.8% | 2.1% |  | 1.9% | 3.8% | 2.8% |
| Not Wild | 1.2% | 2.5% | 1.8% |  | 1.0% | 1.0% | 1.0% |
| Fresh | 2.0% | 1.1% | 1.6% |  | 1.2% | 1.5% | 1.3% |
| Basic/Generic/Blah/Bland/Boring/Packaging | 2.9% | 3.0% | 2.9% |  | 3.7% | 4.9% | 4.3% |
| Packaging/Positive/Clean/Simple/Convenient | 3.0% | 2.6% | 2.8% |  | 4.4% | 3.6% | 4.0% |
| Portion Size/Quantity | 0.8% | 0.7% | 0.8% |  | 2.4% | 2.5% | 2.4% |
| Expensive/High Quality | 0.3% | 0.5% | 0.4% |  | 1.7% | 1.5% | 1.6% |
| Cheap/Inexpensive | 1.2% | 0.5% | 0.8% |  | 0.3% | 1.0% | 0.7% |
| Food/Meal | 0.8% | 1.5% | 1.2% |  | 0.7% | 1.0% | 0.8% |
| Other | 3.7% | 3.4% | 3.6% |  | 4.1% | 3.8% | 3.9% |
| Total | 100% | 100% | 100% |  | 100% | 100% | 100% |

N=1200 (unweighted)

